# Acute administration of a dopamine D2/D3 receptor agonist alters behavioral and neural parameters in adult zebrafish

**DOI:** 10.1101/2022.04.14.488337

**Authors:** Débora Dreher Nabinger, Stefani Altenhofen, Alexis Buatois, Amanda Facciol, Julia Vasconcellos Peixoto, Julia Maria Kuhl da Silva, Gabriel Rübensam, Robert Gerlai, Carla Denise Bonan

## Abstract

The dopaminergic neurotransmitter system is involved in numerous brain functions and behavioral processes. Alterations in this neurotransmitter system are associated with the pathogenesis of several human neurological disorders. Pharmacological agents that interact with the dopaminergic system allow the investigation of dopamine-mediated cellular and molecular responses and may elucidate the biological bases of such disorders. The zebrafish, a translationally relevant biomedical research organism, has been successfully employed in prior psychopharmacology studies. Here, we evaluate the effects of quinpirole (a dopamine D2/D3 receptor agonist) in adult zebrafish on behavioral parameters and neurotransmitter levels. Adult zebrafish received intraperitoneal injections of 0.5, 1.0, or 2.0 mg/kg of quinpirole or saline (control group) twice with an inter-injection interval of 48h. All tests were performed 24h after the second injection. After acute quinpirole administration, zebrafish exhibited decreased locomotor activity, increased anxiety-like behaviors and memory impairment compared to control. However, the quinpirole administration did not affect social and aggressive behavior. Quinpirole-treated fish exhibited altered swimming patterns: fish showed stereotypic swimming characterized by repetitive behavior, swimming from corner to corner at the bottom of the tank preceded and followed by episodes of immobility. Moreover, analysis of neurotransmitter levels in the brain demonstrated a significant increase in glutamate and a decrease in serotonin, while no alterations were observed in dopamine. These findings demonstrate that dopaminergic signaling altered by quinpirole administration results in significant changes in behavior and neurotransmitter levels in the central nervous system of zebrafish. Thus, we conclude that the use of quinpirole administration in adult zebrafish may be an appropriate tool for the analysis of mechanisms underlying neurological disorders related to the dopaminergic system.

## 1 Introduction

The dopaminergic system is involved in several central nervous system (CNS) functions, including the control of movement, reward behaviors, learning, and memory (Beaulieu & Gainetdinov, 2011; Björklund & Dunnett, 2007; Goldman-Rakic, 1997; Jones & Miller, 2008). Alterations of dopaminergic signaling are involved in the pathogenesis of neurodegenerative, neurobehavioral, and psychiatric disorders including Parkinson’s disease, Alzheimer’s disease, Tourette syndrome, schizophrenia, and attention deficit hyperactivity disorder (Armstrong & Okun, 2020; Burns et al., 2019; Chadehumbe et al., 2019; D’Amelio et al., 2018; Howes et al., 2015; Klein et al., 2019). Both pre- and postsynaptic dopamine receptors are present in the nervous system of vertebrates. These G protein-coupled receptors are grouped into two families based upon pharmacological profiles and amino acid sequence similarities: D1-like (D1 and D5) and D2-like (D2, D3, and D4) receptors. Typically, D1-like receptors stimulate adenylyl cyclase activity and produce cyclic adenosine monophosphate (cAMP) as a second messenger, whereas the activation of D2-like receptors inhibits the activation of adenylyl cyclase and also inhibits the production of cAMP and protein kinase A (Beaulieu & Gainetdinov, 2011).

In an attempt to better understand dopamine system functioning and the mechanisms underlying disorders related to this signaling system, receptor agonists and antagonists have been employed. Among them, quinpirole, a psychoactive drug that acts as a selective agonist of dopamine D2 and D3 receptors, has been used to model and investigate mechanisms underlying neurological disorders, including anxiety, schizophrenia, and obsessive-compulsive disorder (Archer & Kostrzewa, 2016; Bortolato & Pittenger, 2017; Brown et al., 2012; Camilla D’Angelo et al., 2014; Stuchlik et al., 2016; Szechtman et al., 2017). In mammals, dopamine D2 receptors are localized both on pre- and postsynaptic dopaminergic neurons, whereas dopamine D3 receptors are localized exclusively postsynaptically (De Mei et al., 2009; Fiorentini et al., 2014). The postsynaptic receptors have an excitatory effect on neurotransmission, while presynaptic receptors act as auto-receptors in a negative feedback loop, inhibiting dopamine release, and consequently presenting an inhibitory effect (Ford, 2014). By acting through these receptors, quinpirole administration leads to endophenotypes associated with some neurological disorders, including changes in locomotor activity, induction of stereotypical responses, elevated erratic and repetitive movements, and changes in learning and memory processes (Abounoori et al., 2020; Bortolato & Pittenger, 2017; Camilla D’Angelo et al., 2014; Irons et al., 2013; Naderi et al., 2016a; b). In addition to mimicking phenotypical features of dopaminergic system-related human CNS disorders, dopamine receptor-selective pharmacological agents may also allow the elucidation of the cellular and molecular bases of these disorders.

The zebrafish has become a popular vertebrate model organism in biomedical research, including neuroscience. Due to its high genetic and physiological homology to mammals, and also because it may be manipulated by various pharmacological and genetic methods, this species has contributed to our understanding of brain function and dysfunction (Gerlai, 2012; Kalueff et al., 2014; Stewart et al., 2014). The dopaminergic system is well characterized in the zebrafish. This neurotransmitter system begins to develop at 15-18 hours post-fertilization (hpf) and by 4 days post-fertilization (dpf), all neuronal cells and their projections are present (Boehmler et al., 2004; Boehmler et al., 2007; Li et al., 2007; Rink & Wullimann, 2001, 2002; Tay et al., 2011). Zebrafish dopamine receptors homologous for all mammalian subtypes have been identified (except for D5), the expression of genes encoding these receptors is detected by 30 hpf, and the receptors themselves are functional by 4 dpf (Boehmler et al., 2004; Boehmler et al., 2007; Li et al., 2007).

The emergence of zebrafish as a valuable model of human CNS diseases has given us insights into the cellular and molecular mechanisms underlying these diseases. Classical neurotransmitter systems involved in several neurological disorders such as dopaminergic, serotoninergic, glutamatergic, and GABAergic are functional in zebrafish (Stewart et al., 2015). Behavioral endophenotypes linked to these disorders, including anxiety-like behavior, stereotypy, impulsivity, decision-making, attention deficit, and depression have also been observed in this animal model (Kalueff et al., 2013). All these characteristics make zebrafish a promising animal model for the study of neurological disorders. We hypothesize that quinpirole exposure may lead to alterations in behavioral patterns, including increased swim activity and anxiety-like behaviors along with alteration in neurotransmitters levels in adult zebrafish. Thus, we investigate the effects of quinpirole administration in adult zebrafish on a variety of behavioral parameters as well as on neurotransmitter levels in the zebrafish brain as a proof-of-concept analysis. We hope that our study sets a precedent and will lead to better understanding of the mechanisms underlying human neurological disorders of the dopaminergic system with the use of zebrafish.

## 2 Materials and Methods

### 2.1 Animals

Adult zebrafish (*Danio rerio*) (5–6 months, 0.2–0.4 g) from wild-type AB background were used. The animals (male and females) were obtained from our breeding colony. Until the treatment, adult zebrafish were maintained in recirculating systems (Zebtec, Tecniplast, Italy) in system water, reverse osmosis-filtered water whose salinity was reconstituted to reach 400–600 μS and pH 6.5–7.5 (ammonia < 0.004 ppm, nitrite < 1 mg/L, nitrate < 50 mg/L, hardness 80–300 mg/L and chloride 0 mg/L). Fish were maintained on 14/10 h light/dark cycle. Fish were fed with paramecium between 6 and 14 days dpf of age, and subsequently received commercial flakes (TetraMin Tropical Flake Fish®) three times a day supplemented with brine shrimp (Westerfield, 2007). All protocols were approved by the Institutional Animal Care Committee from Pontifícia Universidade Católica do Rio Grande do Sul (CEUA-PUCRS, permit number 8181). This study was registered in the Sistema Nacional de Gestão do Patrimônio Genético e Conhecimento Tradicional Associado - SISGEN (Protocol No. A3B073D).

### 2.2 Treatment

Adult zebrafish were exposed to quinpirole by intraperitoneal (i.p.) injection. Quinpirole hydrochloride (Sigma-Aldrich, St. Louis, MO, USA - Q102) was dissolved in water to prepare a stock solution of 1 mg/ml. The test solutions were prepared directly from this stock solution before use. Before each injection, fish were anesthetized with 0.1 g/l tricaine solution (ethyl 3-aminobenzoate methanesulfonate salt). Three different concentrations of quinpirole (0.5, 1.0, and 2.0 mg/kg) were administered, while the control group received the same injection procedure but with saline. A total of two i.p. injections were administered (48h of interval). Each injection consisted of a total volume of 10 μl. Quinpirole concentrations and administration routes were chosen and adjusted based on previous studies (Szechtman et al., 1994a; b; Szechtman et al., 1998) (Fig. 1). The number of injections was established in a pilot experiment. Initially a sequence of injections was tested. The total of two administrations was established because quinpirole had the strongest effect on anxiety-like behavior after this injection regime (Sup. Fig. 1). The behavioral and molecular analyses were performed 24 h after the second injection.

**Fig. 1.**
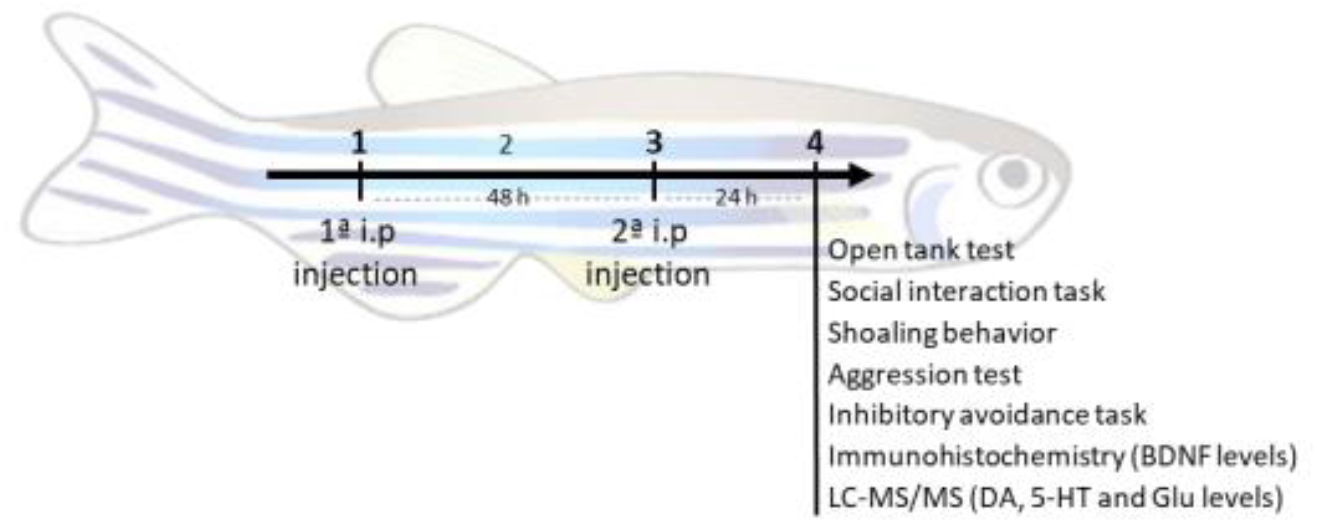
Experimental flowchart. Fish from quinpirole groups received two i.p. injections of the respective quinpirole concentrations (0.5, 1.0, and 2.0 mg/kg) with 48 h of interval. All the tests were evaluated 24 h after the second injection.

### 2.3 Open tank test

Exploratory behavior in the open tank was evaluated 24 h after the second i.p. injection (n = 30 per group). Fish were placed individually in the experimental tanks (30 cm long × 15 cm high × 10 cm wide). After 1 min of familiarization, the swimming path of the fish was recorded for 10 min. The recordings were subsequently analyzed with the EthoVision XT video-tracking software (30 frames per second) (Gerlai et al., 2000; Nabinger et al., 2018). Particular behavioral changes were analyzed using BORIS software, since EthoVision software is not sensitive to detect them. Total distance traveled and time mobile were considered the main parameters of exploration of the novel environment and swim activity. We also measured the absolute turn angle, which quantifies the change in the direction of the center point of the animal between two consecutive video-frames irrespective of the direction of the turn. Absolute turn angle has been found to strongly correlate with erratic movements, and thus may be considered a measure of avoidance response and anxiety (Gerlai et al., 2009). Last, we also quantified the time spent in the bottom zone of the tank, which is also considered an indicator of anxiety-like behavior (Kalueff et al., 2013).

### 2.4 Social behavior

#### 2.4.1 Social interaction task

Social interaction was evaluated 24 h after the last i.p. injection (n = 24 per group). Briefly, fish were individually placed in an experimental tank (30 cm long × 15 cm high × 10 cm wide). On one side of the experimental tank, an empty fish tank was placed; on the other side, a tank containing 15 zebrafish whose size was identical to that of the test fish was presented, the “stimulus tank”. A 5-min session following 1-min familiarization was video recorded for subsequent analysis with EthoVision XT (30 frames per second) (Gerlai et al., 2000). To quantify social interaction and innate preference for conspecifics in detriment of the empty tank, the experimental tank was virtually divided into two halves: a “stimulus zone” closer to the “stimulus tank” and the other remaining half closer to the empty tank. The amount of time the experimental fish spent in the half closer to the stimulus tank was measured.

#### 2.4.2 Shoaling behavior

Shoaling behavior was analyzed by measuring cohesion of the school. After 12-min familiarization, the behavior of groups of five fish (n = 10 shoals, i.e., 50 fish per treatment group) was recorded for 8 min in a test tank (25 cm long × 15 cm high × 25 cm wide). The videos were analyzed using the EthoVision XT software, which tracked the position of each of the five fish in the shoal during the experiment. Using x-y coordinates of individual fish, inter-individual distances between each pair of fish were extracted, and these distances were used as a measure of the shoal cohesion (Buske & Gerlai, 2011; 2014).

### 2.5 Aggression test

The mirror test was used to measure aggressive behavior (n = 24 per group) (Gerlai et al., 2000; Rambo et al., 2017). Each fish was individually placed in an experimental tank (30 cm long × 15 cm high × 10 cm wide). A mirror (45 cm × 38 cm) was placed at the side of the tank at an angle of 22.5° to the back wall of the tank so that the left vertical edge of the mirror touched the side of the tank and the right edge was further away. Thus, when a test fish swam to the left side of the tank, their mirror image appeared closer to them. A 5-min session following 1-min familiarization was video recorded for subsequent quantification of aggressive behavior with EthoVision XT (30 frames per second). The tank was virtually divided into four equal sections allowing counting the number of entries and time spent in each zone. Entry to the zone nearest to the mirror indicated a preference for proximity to the “opponent”. The frequency of entries, the amount of time, and circle swimming movement of the experimental fish spent in each segment were measured for quantification of aggressive behavior.

### 2.6 Inhibitory avoidance task

Inhibitory avoidance task was carried out to assess whether quinpirole administration could impair memory in adult zebrafish (n = 11 per group) (Blank et al., 2009). The task consisted of two sessions, training and test (with 24-h interval between them). In each session, fish were placed individually in an experimental tank (18 cm long × 7 cm high × 9 cm wide), divided by a guillotine door that divided the tank into two equal size, one black (right side) and one white (left side). Throughout the training session, the fish were placed singly in the white compartment with the door closed for 1-min familiarization. After this period, the guillotine door was lifted. Once the fish crossed into the black compartment, the guillotine door was closed, and an electric shock pulse of 3 ± 0.2 V was applied for five seconds. The test subject was then removed from the apparatus and returned to its housing tank for 24 h until the test session. The test session followed the same protocol as the training session’s, except for the electric shock. The latency to enter the black compartment during each session was measured, and the expected increase in the latency at the test session was used as an index of memory retention.

### 2.7 Quantification of neurotransmitter levels

Neurotransmitters were analyzed in zebrafish brain 24 h after the second quinpirole injection, by LC-MS/MS according to Zanandrea et al. (2020). Briefly, animals were euthanized by hypothermal shock and, for each test group, a pool of three brains was collected and homogenized with Ultra Turrax homogenizer (T10 basic IKA®) in 300 μL of ice-cold 0.1 M formic acid (SigmaAldrich, St. Louis, MO) and centrifuged at 20,000g for 20 min at 4 °C. The supernatant was transferred into polypropilene vial insert and injected onto a LC-MS/MS consisted of an LC system (Infinity1290, Agilent Technologies, USA), coupled to a triplequad mass spectrometer (6460 TQQQ, Agilent Technologies, USA). Chromatographic separations were performed on a Zorbax Eclipse Plus C18 RRHD (5 × 2.1 mm, 1.8 μm, Agilent Technologies, USA) column, using a mobile phase consisted of (A) 0.1% formic acid and (B) acetonitrile with 0.1% formic acid, in gradient mode. The gradient started with 66 2% of B, and then programmed to 95% of B after 4.5 min, remaining in this condition for 1.3 min before returning to the start condition. The total chromatographic run was 7.5 min, using a mobile phase flow of 0.2 mL/min, column temperature of 40 °C, and an injection volume of 10 μL. The spectrometer was operated in MRM mode and the analytes were quantified with the following transitions (m/z): dopamine (DA, 154 > 119.1), serotonin (SER, 177 > 160), and glutamic acid (GLU, 148 > 48). Quantifications were performed by external standardization and calibration curves were obtained with the following concentrations: DA and SER 0.1, 0.5, 1.0, 5.0, 10.0, and 50.0 ng/mL; GLU 0.1, 0.5, 0.1, 5.0, 10 and 50.0 μg/mL. Standards were prepared individually in water at concentration of a 0.5 mg/mL and diluted with mobile phase A before LC-MS/MS analysis. The final results were corrected by the protein concentration of each pool and expressed as nanograms (for DA and SER) and micrograms (for GLU) of neurotransmitter per mg of protein. Protein concentrations were determined by the Coomassie blue method (Bradford, 1976), using bovine serum albumin as standard.

### 2.8 Statistical analysis

Nonparametric data were expressed as median ± interquartile range and analyzed by Kruskal-Wallis test followed by a post hoc Dunn’s test (behavioral data). For inhibitory avoidance task, training and test latencies within each group were compared by the Wilcoxon matched-pairs test. The latencies of multiple groups were compared by the Kruskal-Wallis and Mann-Whitney U tests. For parametric statistical tests, normality of data distribution was analyzed by Kolmogorov-Smirnov test. Normally distributed data were expressed as means ± standard error of the mean (S.E.M) and analyzed by one-way ANOVA followed by a post hoc Tukey’s HST test (biochemistry data). For all comparisons, the null hypothesis was rejected when its probability (p) was not more than 5% (p ≤ 0.05).

## 3 Results

### 3.1 Open tank test

The swimming pattern of adult zebrafish was analyzed 24 h after the second quinpirole injection in an open tank test environment. Distance traveled and time mobile were used as the main parameters for exploratory behavior and swimming activity.

Zebrafish injected with quinpirole decreased their activity compared to control (Kruskal-Wallis, H = 34.30; p < 0.0001). A significant decrease in distance travelled in zebrafish injected with 1.0 (p < 0.0001) and 2.0 mg/kg (p < 0.0001) quinpirole doses was observed (Fig. 2a). In addition, a significant decrease in mobility was also observed after quinpirole administration (Kruskal-Wallis, H = 52.58; p < 0.0001). Zebrafish injected with 0.5 (p = 0.001), 1.0 (p < 0.0001) and 2.0 mg/kg (p < 0.0001) exhibited a decrease in time mobile when compared to controls (Fig. 2b).

**Figure 2.**
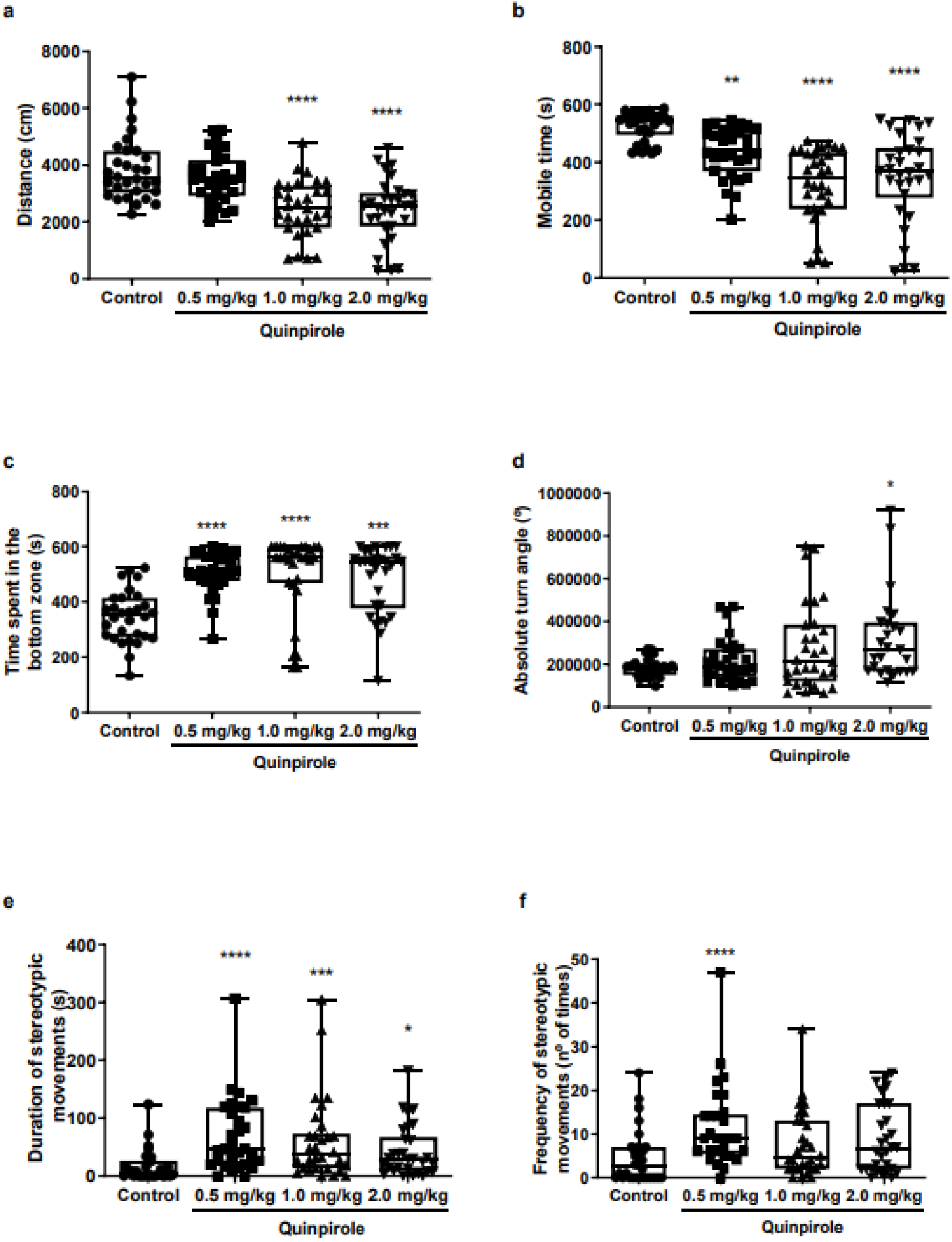
Locomotor activity of adult zebrafish evaluated in the open tank test. Median ± interquartile are shown (each dot represents the individual data). Sample sizes are n = 30 for each group. Distance (a), mobile time (b), time spent in the bottom zone (c), absolute turn angle (d), duration (e) and frequency of erratic movements (f) were analyzed 24 hours after the second i.p. injection. Kruskal-Wallis was used, followed by a post hoc Dunn’s test. Comparisons between control and the quinpirole groups are indicated by asterisk. * indicates significant difference at p ≤ 0.05, ** p ≤ 0.01, *** p ≤ 0.001 and ****p ≤ 0.0001. Note the significant reduction of swimming activity after quinpirole administration (distance (a) and mobile time (b)). Note also that quinpirole administration led to anxiety-like behaviors (time spent in the bottom zone (c), absolute turn angle (d), duration (e) and frequency of erratic movements (f)). For detailed results of statistical analysis, see Results.

To evaluate the potential anxiety altering effect of quinpirole exposure, the time swimming at the bottom of the tank (thigmotaxis) was evaluated. Animals injected with quinpirole presented anxiety-like behavior when compared to controls (Kruskal-Wallis, H = 33.82; p < 0.0001). Fish injected with 0.5 (p < 0.0001), 1.0 (p < 0.0001) and 2.0 mg/kg (p = 0.0001) remained at the bottom of the tank longer than control animals (Fig. 2c). Furthermore, quinpirole treated fish exhibited increased absolute turn angle, likely an indication of elevated erratic movements, a sign of increased fear or anxiety (Kruskal-Wallis, H = 8.290, p = 0.0404). Fish injected with the highest concentration tested (2.0 mg/kg, p = 0.0314) exhibited increased turn angle when compared to controls (Fig. 2d).

Additionally, animals exposed to quinpirole presented an altered swimming pattern (Kruskal-Wallis, H = 24.26; p = 0.0404). After quinpirole injections, more than 90% of the animals injected with 0.5 (p < 0.0001), 1.0 (p = 0.0015) and 2.0 mg/kg (p = 0.0262) presented a stereotypic swimming characterized by the pattern of rigid and repetitive behavior. The fish swam from corner to corner at the bottom of the tank multiple times with high velocity; the changes of direction during this stereotypic swimming were made by rapid turning by a single bout of movement; the sequence of repetitive stereotypic movements was preceded and followed by immobility episodes. All fish injected with quinpirole that performed this behavior presented this same pattern, with time and number of episodes of repetitions varying among the groups (Fig. 2e, f).

### 3.2 Social behavior

There were no alterations in social behavior after quinpirole injections. Zebrafish injected with quinpirole presented the same preference for the stimulus area as controls when evaluated individually at the social interaction task (Kruskal-Wallis, Time in the stimulus zone: H = 1.476, p = 0.6879; Frequency in the stimulus zone: H = 4.448, p = 0.2170) (Fig. 3). Furthermore, no alterations in the shoal cohesion were observed when fish were tested in groups in the shoaling behavior task (Kruskal-Wallis, Distance between subjects: H = 9.800, p = 0.203; Proximity: H = 5.752; p = 0.1243; Body contact: H = 2.333, p = 0.5063) (Fig. 4).

**Figure 3.**
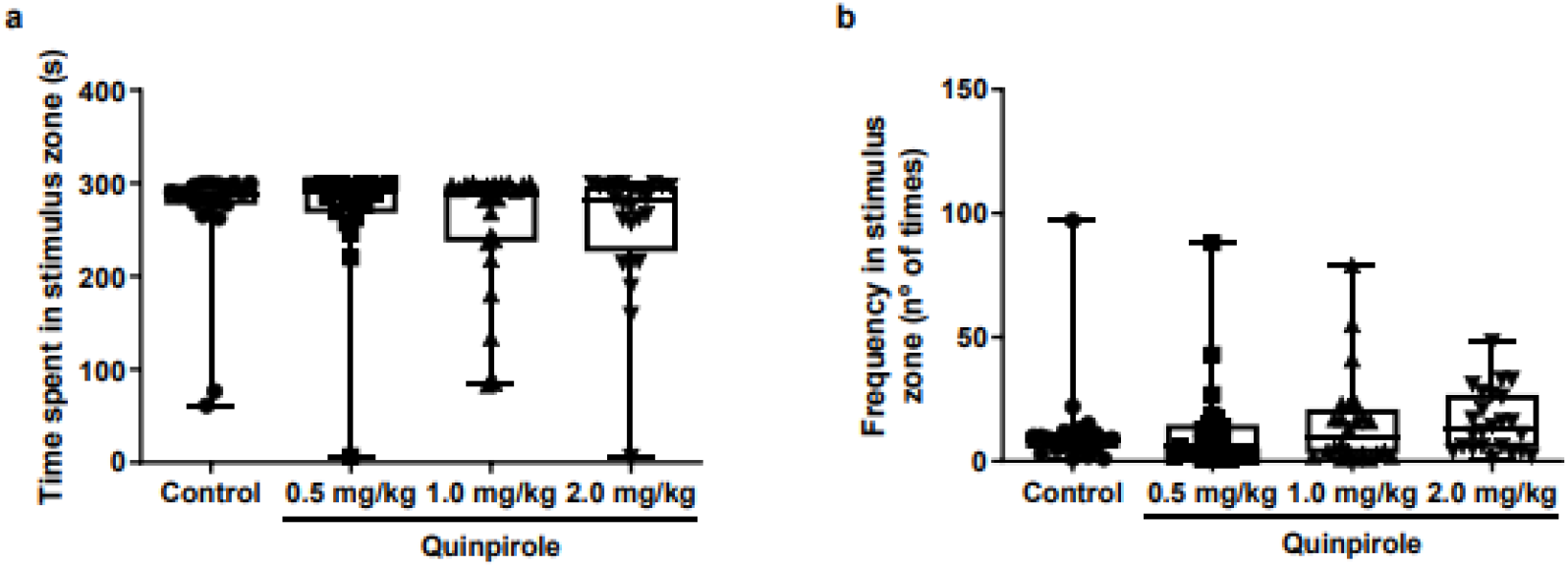
Analysis of social behavior in adult zebrafish individually tested at the social interaction task. Median ± interquartile are shown (each dot represents the individual data). Sample sizes are n = 20 for control group and n = 24 for quinpirole groups. Time spent in stimulus zone (a) and frequency in stimulus zone (b) were analyzed 24 hours after the second i.p. injection. Kruskal-Wallis was used, followed by a post hoc Dunn’s test. Zebrafish injected with quinpirole presented the same preference for the stimulus area as controls when evaluated individually at social interaction task. For detailed results of statistical analyses, see Results.

**Figure 4.**
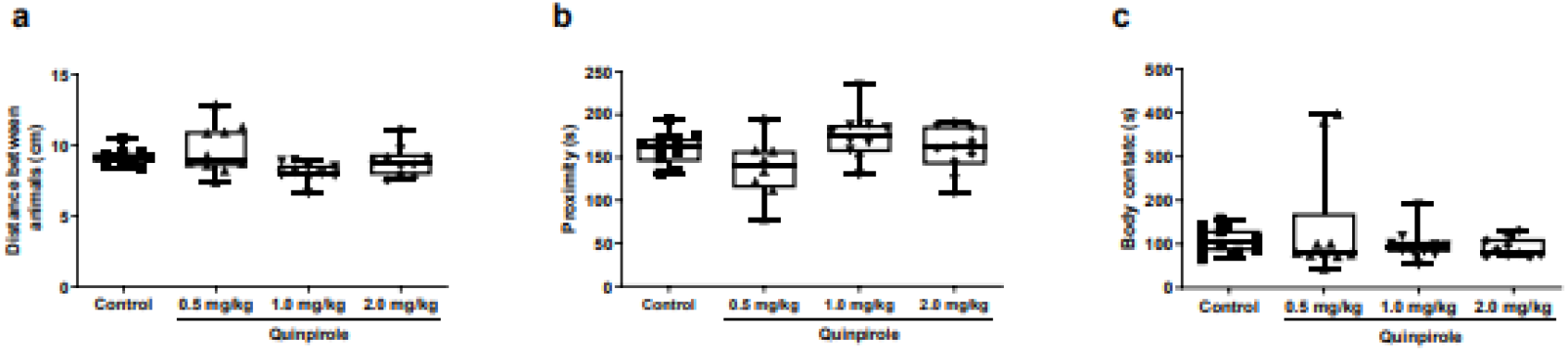
Analysis of social behavior in adult zebrafish evaluated in groups (shoaling behavior). Median ± interquartile are shown (each dot represents the individual data). Sample sizes are n = 10 for each group (5 fish per trial, 50 fish per group). Distance between animals (a), proximity (b) and body contact (c) were analyzed 24 hours after the second i.p. injection. Kruskal-Wallis was used, followed by a post hoc Dunn’s test. No alterations in the school cohesion were observed when fish were tested in groups at shoaling behavior analysis. For detailed results of statistical analyses, see Results.

### 3.3 Aggression test

The quinpirole administration did not induce any alteration in aggressive behavior at any doses employed. Zebrafish injected with quinpirole presented the same response to the mirror as control fish (Kruskal-Wallis, Time in the zone nearest to the mirror: H = 1.050, p = 0.7892; Frequency in the zone nearest to the mirror: H = 3.081, p = 0.3792; Rotation: H = 5.814, p = 0.1210) (Fig. 5).

**Figure 5.**
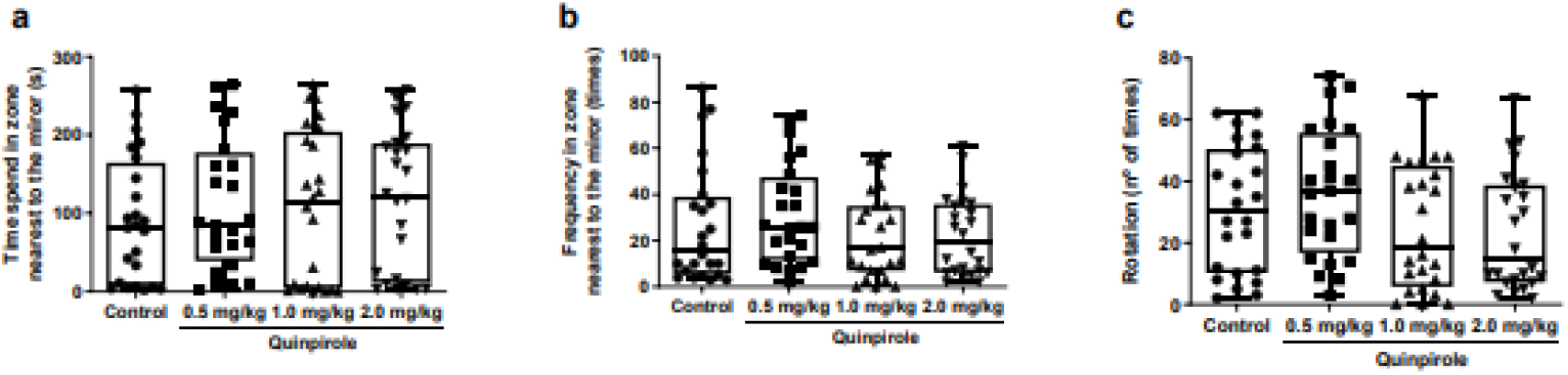
Evaluation of aggressive behavior in adult zebrafish in the mirror-induced aggression task. Median ± interquartile are shown (each dot represents the individual data). Sample sizes are n = 24 for each group. Time spent in the zone nearest to the mirror (a), frequency in the zone nearest to the mirror (b) and frequency of rotations (c) were analyzed 24 hours after the second i.p. injection. Kruskal-Wallis was used, followed by a post hoc Dunn’s test. No alterations were observed after quinpirole administration on the parameters evaluated in the mirror-induced aggression task. For detailed results of statistical analyses, see Results.

### 3.4 Inhibitory avoidance task

Memory was evaluated by the inhibitory avoidance task. Zebrafish injected with quinpirole (0.5, 1.0, and 2.0 mg/kg) exhibited a quasi-linear dose dependent memory performance impairment, with the highest dose group showing the most and the lowest dose the least robust impairment. There was no difference in the latency to enter the dark compartment between training and test sessions in zebrafish of any of the quinpirole groups, indicating an impairment of aversive memory, whereas there was a significant difference in the control group (p < 0.0001) (Fig. 6a). Comparing the groups only at the test session, there is a significant decreasing latency at fish injected with 1.0 (Mann-Whitney, W = 9.5; p = 0.01) and 2.0 mg/kg (Mann-Whitney, W = 2; p = 0.001) when compared to controls (Fig. 6b).

**Figure 6.**
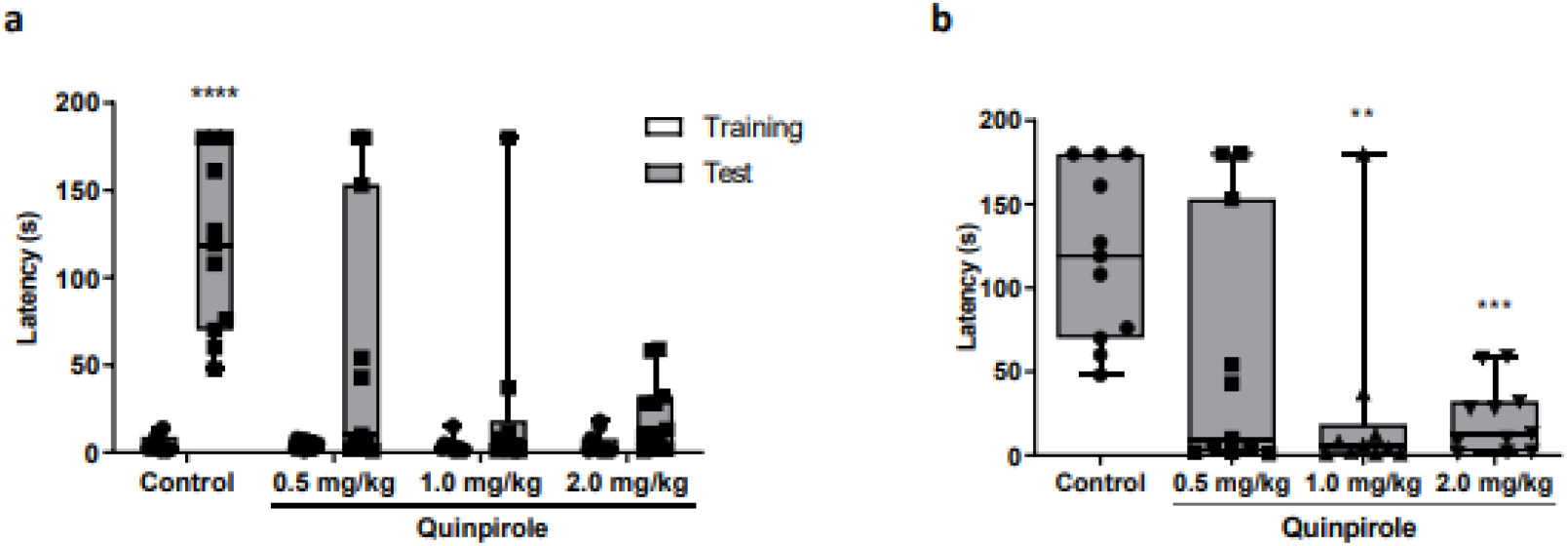
Inhibitory avoidance task. Sample sizes are n = 11 for each group. Training and test sessions are indicated by white and grey bars, respectively. Median ± interquartile are shown (each dot represents the individual data). Inhibitory avoidance training and test latencies within each group were compared by the Wilcoxon matched pairs test. Latencies of multiple groups were compared by the Kruskal-Wallis and Mann-Whitney U tests (a). Latencies between groups at training session were compared by Mann-Whitney (b). Latencies differences between training and test session are indicated by asterisk. ** indicate significant difference at p ≤ 0.01, *** p ≤ 0.001 **** p ≤ 0.0001. Note that quinpirole administration caused memory impairment. Only control animals presented an increased latency to enter the dark compartment during test session when compared to training session. For detailed results of statistical analyses, see Results.

### 3.5 Quantification of neurotransmitter levels

Brain levels of dopamine, serotonin, and glutamate were evaluated 24 h after the second injection in zebrafish (Fig. 7). ANOVA found no significant change in Dopamine (F_(3, 19)_ = 1.245, p = 0.3213), but revealed a significant treatment effect for Glutamate (F_(3, 19)_ = 6.281, p = 0.0038) and Serotonin (F_(3, 19)_ = 5.236, p = 0.0084). Subsequent Tukey HSD post hoc multiple comparison test showed that fish injected with 2.0 mg/kg exhibited a significant (p = 0.0311) increase in glutamate levels. In addition, a significant decrease in serotonin levels was observed in fish injected with 0.5 (p = 0.0188), 1.0 (p = 0.0193) and 2.0 mg/kg (p = 0.0155) quinpirole.

**Figure 7.**
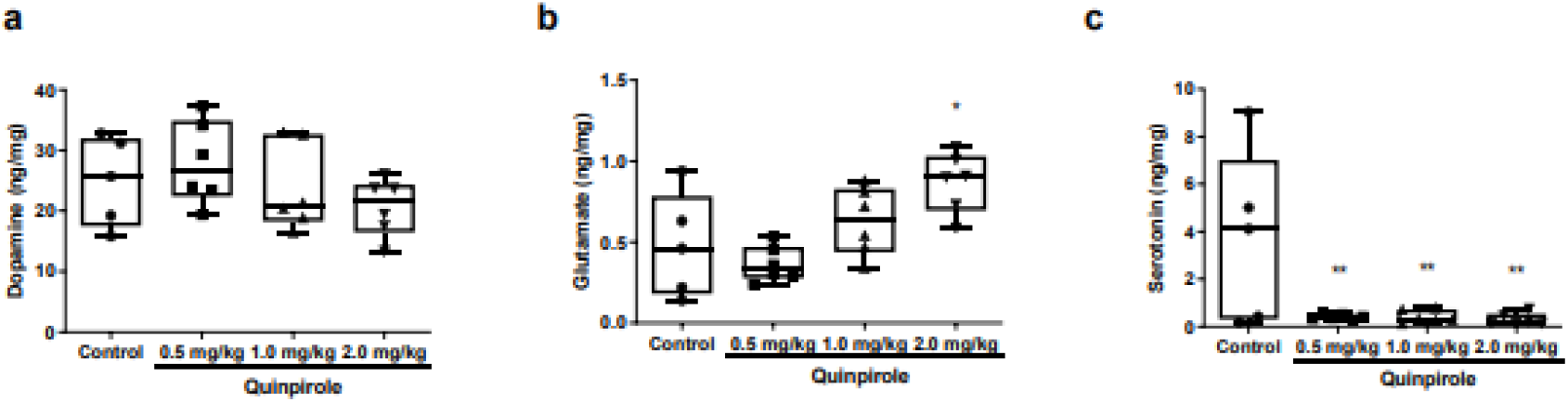
Levels of dopamine (a), glutamate (b) and serotonin (c) in the adult zebrafish brain after quinpirole administration. Median ± interquartile are shown (each dot represents the individual data). Sample sizes are n = 6 for each group (pool of three brains for each sample). One-way ANOVA was used, followed by post-hoc Tukey’s test. Comparisons between control and the quinpirole groups are indicated by asterisk. * indicates significant difference at p ≤ 0.05 and ** p ≤ 0.01. Note significant increase in glutamate level after quinpirole administration at dose of 2.0 mg/kg and the significance reduction of serotonin at all quinpirole doses tested. For detailed results of statistical analyses, see Results.

## 4 Discussion

In the present study, we investigated the effects of acute quinpirole administration in adult zebrafish on behavioral parameters and neurotransmitter levels. After acute quinpirole administration, fish exhibited decreased locomotor activity, increased anxiety-like behavior, including elevated erratic movements, as well as learning/memory performance impairment when compared to controls. On the other hand, we found quinpirole administration not to affect social and aggressive behaviors. Also notable is the change we observed in the swimming pattern of fish treated with quinpirole. These fish showed a stereotypic swimming pattern characterized by repetitive behavior, swimming from corner to corner at the bottom of the tank preceded and followed by episodes of immobility. Moreover, after quinpirole injections, quantification of neurotransmitter levels showed a significant increase in glutamate and a decrease in serotonin, while no alteration was observed in dopamine levels.

Dopamine is an important modulator of locomotor activity in mammals. In zebrafish, this neurotransmitter also regulates the motor activity and is required for the initiation of movement (Ek et al., 2016; Irons et al., 2013; Lambert et al., 2012; Souza et al., 2011; Thirumalai & Cline, 2008). In this study, quinpirole was found to decrease distance traveled and mobile time at all doses tested. Results regarding the effects of quinpirole exposure on locomotor activity have been controversial in zebrafish studies. Quinpirole exposure in early life stages of zebrafish (around 5 and 7 dpf) has been found to increase as well as decrease locomotor activity. Boehmler et al. (2007) observed hyperactivity after 1-hour exposure at a concentration of 12.5 μM quinpirole, a finding consistent with previous reports. Using similar concentrations and length of exposure, Irons et al. (2013) described increased locomotor activity in a light/dark task. In the dark, quinpirole increased activity at 16.7 μM whereas hyperactivity was seen at 5.5 and 16.7 μM quinpirole concentrations in the light. In contrast, two other studies found decreased movements and activity after exposure to 10 and 30 μM quinpirole (Lange et al., 2018; Souza et al., 2011). Other studies have shown no effects of quinpirole on locomotor activity after exposure to 1 mg/l concentration of this drug in adult zebrafish (Naderi et al., 2016a; b).

The contradicting findings may be due to different procedures and/or genetic background of fish used, and to the fact that quinpirole may have a dose-dependent biphasic profile, a possibility already confirmed in mammals, with decreased activity induced by low concentrations and increased activity by high concentrations (Li et al., 2010). Such biphasic dose-response is assumed to result from the dose-dependent activation of the D2 versus D3 receptor (De Mei et al., 2009). The D2 receptor is expressed in several brain regions both in the pre- and postsynaptic terminal of synapses whereas the D3 receptor is expressed exclusively postsynaptically in the brain of mammals in several brain areas (De Mei et al., 2009; Fiorentini et al., 2014). In mammals, low concentrations of D2 receptor agonists have been found to primarily activate presynaptic receptors, which serve as auto-receptors in a negative feedback loop, reducing dopamine release and decreasing locomotion. At high concentrations, D2 receptor agonists induce hyperactivity due to the agonists’ action on the postsynaptically localized receptor (Beaulieu & Gainetdinov, 2011; Beninger et al., 1991; Fiorentini et al., 2014; Li et al., 2010; Missale et al., 1998; Van der Weide et al., 1988). In zebrafish, the dopamine D2 and D3 receptors are found both pre- and postsynaptically. In the first hours of development, the receptors are localized in the postsynaptic membrane. However, as development progresses, expression of D2 and D3 receptor genes extends, and the microstructural localization changes from exclusively postsynaptic to both the pre- (auto-receptors) and the postsynaptic membrane of neurons (Boehmler et al., 2004).

In addition to regulating locomotor activity, dopamine, just like in mammals, is also involved in a variety of neurobehavioral functions and phenomena in zebrafish, including anxiety, aggression, social behavior, and learning and memory (Irons et al., 2013; Kacprzak et al., 2017; Liu et al., 2020; Naderi et al., 2016a; b; Scerbina et al., 2012; Teles et al., 2013). We found quinpirole administration to elevate anxiety-like behavior and induce stereotypic swimming movements, characterized by repetitive behavior, but we detected no effect of this treatment on social and aggressive behaviors.

To the best of our knowledge, such phenotypes have not been demonstrated in zebrafish after quinpirole exposure. Other dopamine agonists and partial agonists of D2 and D3 dopamine receptors have been administered to zebrafish and were found to have different effects. For example, the administration of partial agonists of D2 receptors such as aripiprazole and modafinil has been shown to have an anxiolytic effect in adult zebrafish (Barcellos et al., 2020; Johnson & Hamilton, 2017). Another partial agonist, ketamine, has been found to both increase and decrease anxiety in adult zebrafish, contradictory findings (De Campos et al., 2015; Pittman & Hylton, 2015; Riehl et al., 2011). Furthermore, during early life stages of development, this partial agonist of the D2 dopamine receptor was found to induce anxiety-like behavior in zebrafish larvae (Félix, et al., 2017a; b). Besides the effects related to anxiety, ketamine has also been found to induce stereotypical behaviors, evoke circular swimming, reduce aggression, and disrupt shoaling in zebrafish (Michelotti et al., 2018; Riehl et al., 2011; Zakhary et al., 2011). Apomorphine, a non-selective agonist of the D3 dopamine receptor, induced anxiolytic effects in zebrafish larvae (Ek et al., 2016). In the same way, partial agonists as aripiprazole and busperidone also decreased anxiety (Barcellos et al., 2020; Müller et al., 2020).

Some of our results are in line with what was observed in these previous studies. However, as mentioned above, these studies used partial agonists of D2 and D3 dopamine receptors. Partial agonists also act via other receptors from different signaling systems, which perhaps explains why some of these prior results differ from those we present here. Although the effects observed in our study are not completely consistent with other studies in zebrafish using other D2 and D3 dopamine receptor agonists, our findings agree with observations obtained with rodents after quinpirole exposure. Quinpirole administration has been used in rats and mice to model psychiatric disorders, such as obsessive-compulsive disorder and schizophrenia. After exposure, the animals presented anxiety-like behavior and elevated stereotypic and erratic movements (Hoffman, 2011; Kostrzewa et al., 2016; Navarro & Maldonado, 1999; Nielsen et al., 2017; Sams-Dodd, 1998; Szechtman et al., 2017). Additionally, only a few studies have reported alterations in social and aggressive behavior (Maple et al., 2017; Sams-Dodd, 1998; Szechtman et al., 2017).

The dopaminergic neurotransmitter system plays a role in learning and memory processes (Goldman-Rakic, 1997). In zebrafish, exposure to the same concentration of quinpirole resulted in contradictory results in different learning and memory tasks. Using a plus-maze to assess associative learning performance, activation of D2/D3 receptors has been found to enhance memory acquisition, while their blockade has been found to impair learning performance (Naderi et al., 2016a). The same group, using a latent learning paradigm to assess cognitive performance, observed that quinpirole significantly impaired memory acquisition and consolidation in zebrafish, and the antagonism of D2 receptors enhanced latent learning ability (Naderi et al., 2016b). Similar to zebrafish, rodent studies also show controversial effects of quinpirole on learning and memory performance. In a fear conditioning paradigm, the administration of quinpirole was found either to improve or to impair learning and memory processes in rodents (Farahmandfar et al., 2016; Lénárd et al., 2017; Nader & LeDoux, 1999a;b). In the present study, using the inhibitory avoidance task, quinpirole administration was found to impair learning/memory performance in adult zebrafish in a dose-dependent manner. To the best of our knowledge, this is the first demonstration of quinpirole effects in inhibitory avoidance learning/memory performance in zebrafish. Among the modulatory systems involved in the regulation of information acquisition and synaptic plasticity in the CNS, the dopaminergic system has received particular attention (Wise, 2004). It is not surprising that dysfunction in dopaminergic neurotransmission is associated with a variety of psychiatric and neurodegenerative disorders in which memory impairment is an important symptom (Armstrong & Okun, 2020; D’Amelio et al., 2018; Howes et al., 2015; Klein et al., 2019). In this regard, a growing body of evidence indicates an involvement of dopamine receptors in acquisition, consolidation, retrieval, reconsolidation, and extinction phases of different types of learning and memory (El-Ghundi et al., 2007; Puig et al., 2014). Previous studies identified a differential involvement of D2 receptor signaling in the acquisition of certain types of memory (Brown et al., 2000; Kurylo et al., 2004; Merritt & Bachtell, 2013; Ponnusamy et al., 2005; Yawata et al., 2012; Young et al., 2014), and this may be one of the reasons for the contradictory findings. Also, as mentioned, depending on the quinpirole concentration pre- or postsynaptic D2/D3 receptors can be activated. It has been demonstrated that activation of postsynaptic receptors led to recall of memory (Bracs et al., 1984; Ichihara et al., 1988; Naderi et al., 2016a); likely this could indicate that the impaired memory observed here results from presynaptic activation of D2 receptors. Further studies are needed to elucidate the role of D2/D3 receptors in zebrafish memory processes.

Considering that quinpirole exposure by itself alters the neurotransmitter systems, we decided to evaluate whether quinpirole exposure could impair neurotransmitters levels. The determination of neurotransmitter levels in the overall zebrafish brain after quinpirole administration showed a significant increase in glutamate and a decrease in serotonin levels, while no alterations were observed in dopamine levels. To the best of our knowledge, the assessment of neurotransmitter levels in the zebrafish brain after quinpirole exposure has not been previously performed. In rodents, such analysis has been attempted, but the observed results vary according to the brain area analyzed. For example, after quinpirole administration, glutamate is reported to have decreased levels in nucleus accumbens, globus pallidus, and striatum, and increased levels in the nucleus accumbens and substantia nigra (Abarca et al., 1995; Beggiato et al., 2016; Biggs et al., 1997; Dalia et al., 1998; Escobar et al., 2015; Ferraro et al., 2012a; b; Kalivas et al., 1997; Krügel et al., 2004; Tanganelli et al., 2004). Dopamine has been shown to have decreased levels in the nucleus accumbens and prefrontal cortex, and when evaluated in the whole brain, the overall level of dopamine was found to be increased (Brodnik et al., 2013; Chen et al., 1987; De Haas et al., 2011; Escobar et al., 2015; 2017; Ferré & Artigas, 1995; Koeltzow et al., 2003; Liu & Steketee, 2011; Sullivan et al., 1998; Thorré et al., 1998). For serotonin, an increased level in the dorsal raphe and decreased level in the substantia nigra was observed, while no alterations were found in the nucleus accumbens (Ferré & Artigas, 1995; Koeltzow et al., 2003; Martín-Ruiz et al., 2001; Thorré et al., 1998). Although quinpirole is a dopamine receptor selective drug, the most robust changes in neurotransmitter levels we found were in glutamate (increased) and in serotonin (decreased) but not in dopamine. While this may seem surprising, it actually is in line with a large body of data, showing intricate interaction among neurotransmitter systems. Nevertheless, at this point, we do not know via what pathways/circuits exactly these changes may have been induced by quinpirole in the zebrafish brain.

Alterations in neurotransmitter levels are known to affect behavior, and some of the ones reported here are in agreement with what was reported before. Although we did not observe a statistically significant decrease in dopamine levels, the results showed a downward trend. In the literature, reduced dopamine levels in the zebrafish brain have been correlated with decreased swim activity, impulsive-like behavior and increased anxiety-like behavior (Bortolotto et al., 2014; Herculano & Maximino, 2014; Huang et al., 2015; Kyzar et al., 2013; Shams et al., 2018). Furthermore, decreased shoal cohesion, disrupted social preference, and impaired acquisition and consolidation of memory were also reported in zebrafish that presented low levels of dopamine (Bortolotto et al., 2014; Scerbina et al., 2012; Shams et al., 2018). Similarly to what we observed in the present study, zebrafish with increased glutamate levels exhibited abnormal tracking pattern and anxiety-like behavior (Kundap et al., 2017). Furthermore, reduced glutamate reuptake was observed in animals that displayed lower total distance traveled and swimming speed, and increased time of immobility (Mussulini et al., 2018). In addition, administration of NMDA was found to increase freezing and erratic movements (Herculano et al., 2015). Last, decreased serotonin levels were observed in animals with increased anxiety-like behavior (Kyzar et al., 2013; Herculano & Maximino, 2014; Maximino et al., 2013).

These neurotransmitters are known to be implicated in the pathogenesis of many neuropsychiatric disorders. For example, perturbed glutamatergic signaling has been implicated in the pathogenesis of anxiety disorders, depression, bipolar disorder, and schizophrenia (Mattson, 2008). Disturbances in serotonin signaling are involved in obsessive– compulsive disorder, bulimia, chronic impulsivity, aggression, and suicide (Martinowich & Lu, 2008). Last, dopamine is known to be involved in the pathogenesis of Parkinson’s disease, Alzheimer’s disease, Tourette syndrome, schizophrenia, and attention deficit hyperactivity disorder (Klein et al., 2019). The altered behaviors reported here, and in the previous studies, in combination with the altered signaling of neurotransmitters related to such psychiatric disorders, reinforce the importance and relevance of the use of animal models in the analysis of these human central nervous system disorders. Thus, in summary, our study shows that acute quinpirole administration affects adult zebrafish behavior and alters brain levels of serotonin and glutamate in the adult zebrafish. The observed alterations demonstrate the involvement of the dopaminergic neurotransmitter system in a variety of brain functions and behavior and show that the zebrafish is psychopharmacologically similar to mammalian model organisms. Thus, our proof-of-concept study opens a new avenue to the mechanistic analysis of the dopaminergic system and the modeling of neuropsychiatric disorders using this simple vertebrate.

## Supplementary figure

**Supplementary figure 1.**
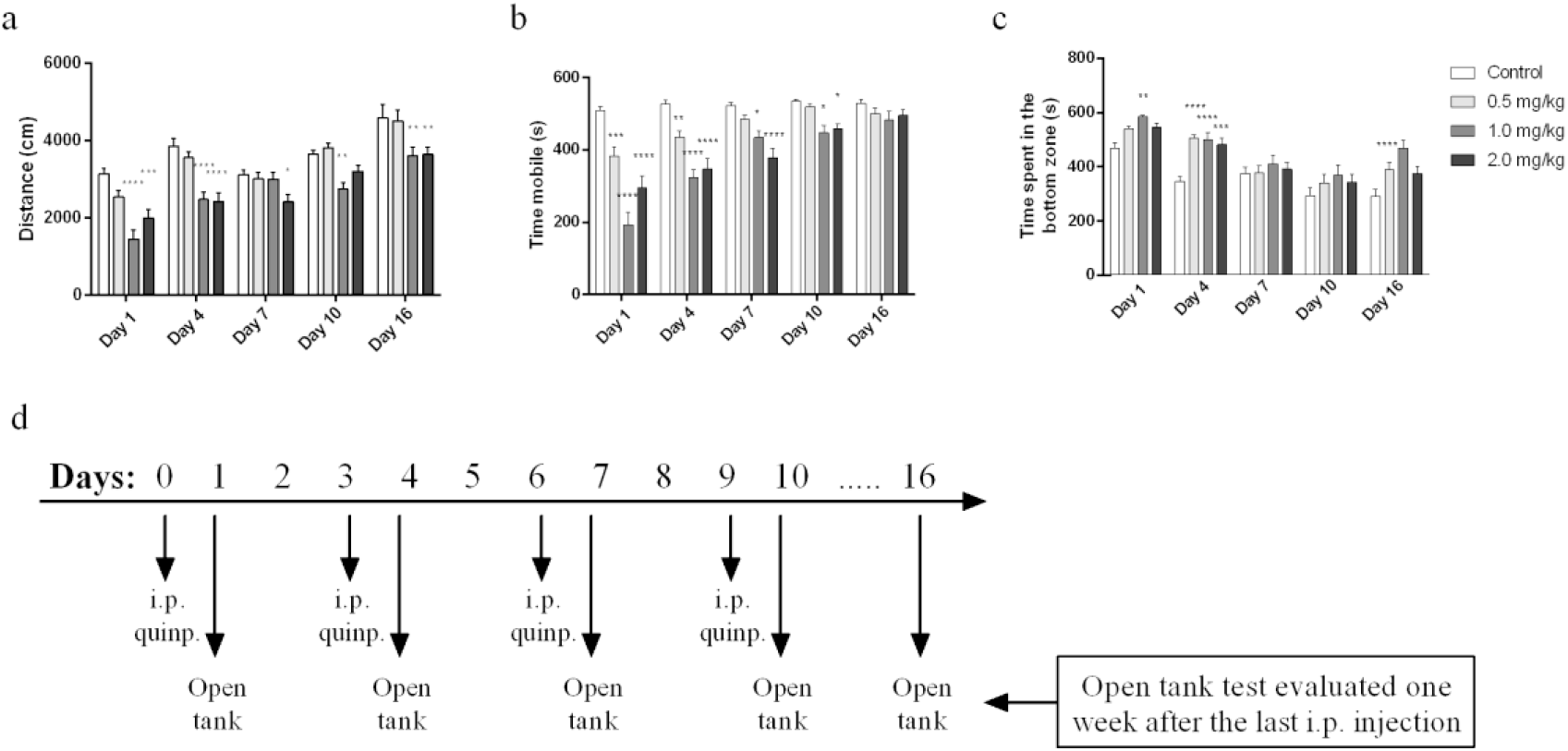
Locomotor activity of adult zebrafish evaluated in the open tank test. Mean ± SEM are shown. Sample sizes are n = 30 for each group. Distance (a), mobile time (b), time spent in the bottom zone (c) were analyzed 24 hours after each i.p. injection. (d) protocol of injections and behavior analysis. Two-way ANOVA was used, followed by a post hoc Bonferroni test. Comparisons between control and the quinpirole groups are indicated by asterisk. * indicates significant difference at p ≤ 0.05, ** p ≤ 0.01, *** p ≤ 0.001 and **** p ≤ 0.0001.

